# Assembly of branched chain amino acid (BCAA) to toxic fibrils may be related to pathogenesis of Maple syrup urine disease (MSUD)

**DOI:** 10.1101/2024.10.01.616096

**Authors:** Chandra P Kanth, Monisha Patel, Raj Dave, Ankur Singh, Aayushi Joshi, Manoj Kumar Pandey, Dhiraj Bhatia, Nidhi Gour

**Affiliations:** Department of Chemistry, Pandit Deendayal Petroleum University, Gujarat, India; School of Science, Indrashil University, Kadi, Mehsana, Gujarat, 382740, India; Department of Biological Engineering Discipline, Indian Institute of Technology, Palaj, Gujarat, 382355, India

**Keywords:** Inborn errors of metabolism, branched chain amino acid, generic amyloid hypothesis, self-assembly

## Abstract

Inborn errors of metabolism (IEMs) are a group of diseases caused by mutations in single genes, leading to the buildup of metabolites that are typically toxic or disrupt normal cellular function. The etiological relation of metabolic disorders has been uncovered through the study of metabolite amyloids. Various metabolites that accumulate in IEMs have been reported to self-assemble into organized structures. These structures exhibit similar physicochemical properties as proteinaceous amyloid fibrils. In this context, our study illustrated the aggregation properties of Branched chain amino acid (BCAA) i.e. Isoleucine, Leucine and Valine that accumulate in Maple syrup urine disease (MSUD) to investigate their propensities to assemble into amyloid-like fibrils. The structural morphologies of BCAA were studied via. microscopic techniques like Scanning electron microscopy (SEM), optical microscopy and phase contrast microscopy. Further, characterization techniques were employed to understand the physicochemical properties of the self-assemblies and its underlying mechanism. The amyloid-like nature of these aggregates was confirmed using Thioflavin T (ThT) and Congo Red (CR) assays, indicating a possible cytotoxic effect. The MTT assay reveals BCAAs were cytotoxic and significantly decrease cell viability. Our study plays a key role in understanding the physicochemical properties of MSUD in association to amyloid disease, possibly paving the way for the development of therapeutic solutions in the future.

**Graphical Abstract:** 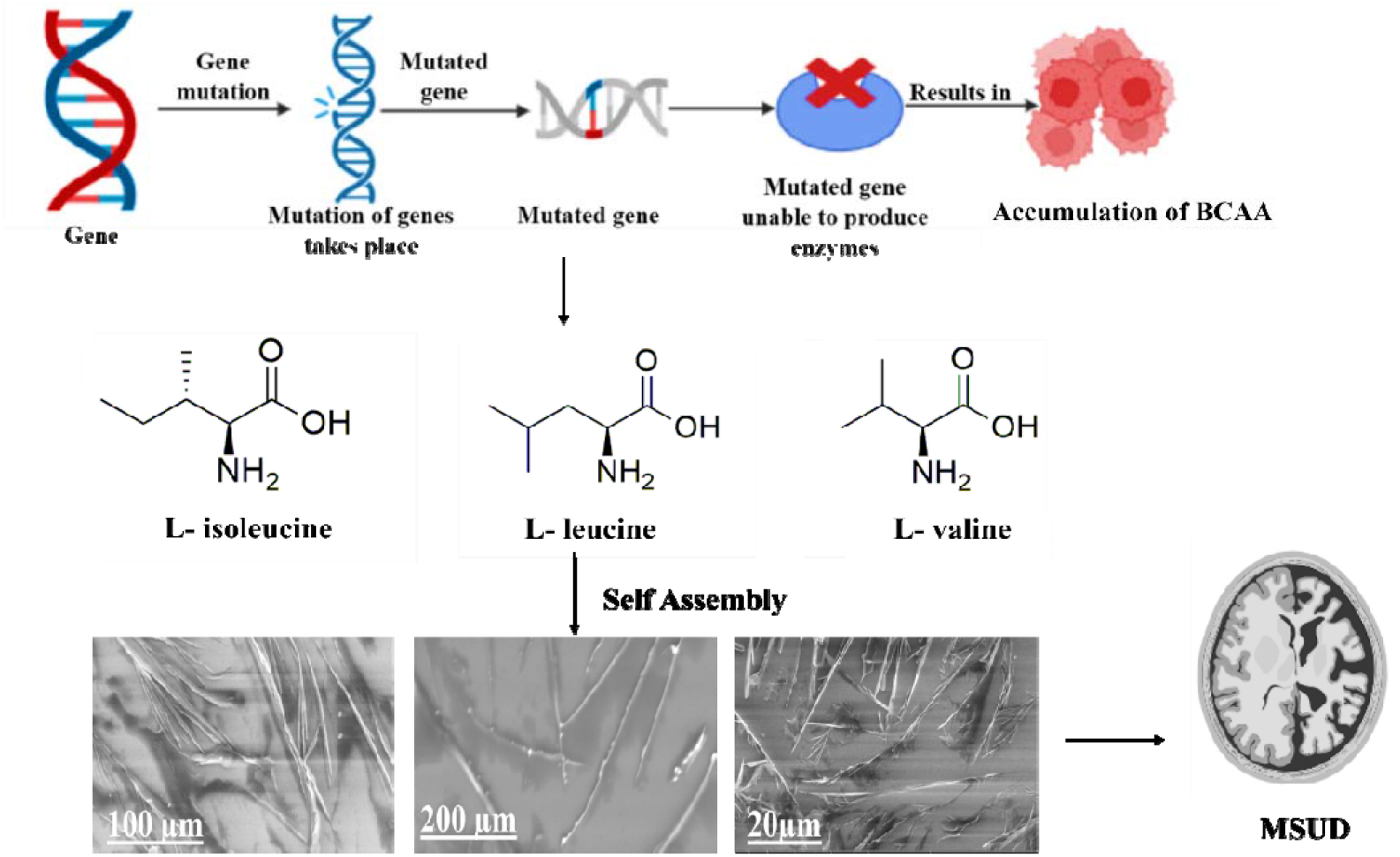

The self-assembly of BCAA-Ile, Leu, Val was investigated and the findings indicate that their aggregation may follow an amyloidogenic pathway.

## Introduction

Maple syrup urine disease (MSUD) is an inborn error of metabolism which is caused by the deficiency of branched chain α-ketoacid dehydrogenase complex enzyme, which causes branched chain amino acid (BCAA) catabolism.^1,2^ MSUD hence leads to accumulation of BCAAs isoleucine, leucine and valine and its elevated levels are found in the blood plasma, cerebrospinal fluid and urine.^3,4^ The patho-physiology of MSUD is accompanied by the acute neurological deterioration and severe encephalopathies in the brain.^5,6^ Although, the exact mechanism by which MSUD causes neuropathology and brain damage are still poorly understood, literature reports suggest that MSUD is associated with the reduced mitochondrial functioning and increased oxidative stress which induces cell death by apoptosis.^7-10^ Literature reports also reveal that when the concentration of MSUD metabolites like BCAA are increased in the culture, it causes neurotoxic effects and induces apoptosis in glial and neuronal cells.^11, 12^ Further, it has been shown that chronic subcutaneous administration of BCAA in mouse models induce long term memory loss and impairment of spatial memory.^13^ Hence, the pathologies associated with MSUD exhibit striking resemblance to other amyloid associated diseases like Alzheimer’s and Parkinson’s which cause neurodegeneration and memory loss and indicative that MSUD might have etiological relation to aggregation associated diseases.^14,15^ Recently, a report by Gazit and coworkers have also shown the self-assembly of BCAA which suggests a close relation between MSUD and amyloids.^16^

The self-assembly of phenylalanine (Phe) to amyloid like toxic fibers and its crucial significance in the etiology of Phenylketonuria was the first ground breaking report which suggested a close association between IEMs and amyloid related diseases.^17,18^ Further a “generic amyloid hypothesis” was proposed wherein Gazit and coworkers suggested that various inborn error of metabolisms (IEM’s) might have a common origin and the metabolites might be assembling to amyloid-like toxic fibers which leads to the pathogenesis of metabolic diseases.^17^ In context of this hypothesis, they further reported amyloid-like structures formed by the self-assembly of non-proteinaceous metabolites like adenine, orotic acid, uridine and cystine.^19^ Further, the same research group reported formation of amyloid-like fibers by self-assembly of tryptophan (Trp)^20^ and tyrosine (Tyr)^21^ and attributed pathology of diseases like hypertryptophanemia and tyrosinuria to these toxic assemblies. Our group is also been interested in studying the self-assembly of metabolites and investigating its association in the etiology in IEMs. In this context, we have reported the self-assembly of single amino acids and metabolites. Our studies suggest amino acids like cysteine, methionine, proline, hydroxyproline lysine, glutamine, glutamic acid, aspartic acid assembles to amyloid-like structures.^22-24^ Further, metabolites of urea and uric acid cycle, homogentisic acid, isovaleric acid and N-acetyl aspartic acid also reveals similar propensities towards amyloid like structure formation suggesting implications of their self-assembly in the pathogenesis of IEMs in excess of these metabolites.^25,26^ Hence the studies conducted so far on single amino acid assemblies are in accordance with “generic amyloid hypothesis” and suggest the underlying cause of pathogenesis in variety of single amino acid metabolic disorders might be attributed to toxic amyloid like assemblies with the IEMs.

However, this studies were mainly explored in biological perspective of BCAA assemblies. The intermolecular forces which leads to BCAA assemblies were not studied from a chemical perspective. Hence, motivated by our own previous work and the recent research wherein MSUD has been studied in the context with BCAA assemblies, we were motivated in studying the self-assembly behavior via different biophysical assays to access the role of intermolecular forces in causing their self-assembly. Interestingly, our studies suggest that BCAA, indeed assemble to amyloid like toxic fibers. The amyloid like fibrillar morphologies of BCAA were studied by microscopic techniques like Scanning electron microscopy (SEM), optical microscopy and phase contrast microscopy. The physicochemical properties of the self-assemblies and its underlying mechanism were further studied by various solid state characterization techniques like solid state nuclear magnetic resonance (NMR), fourier-transform infrared (FTIR), thermogravimetric analysis (TGA) and X-ray diffraction (XRD). The amyloid nature of the assemblies was deciphered by assessing the interaction between BCAA fibers and amyloid specific dyes Thioflavin T (ThT) and Congo Red (CR). Finally, the cytotoxicity analysis was done by MTT assay which revealed all three Ile, Leu and Val were cytotoxic and reduced cell viability significantly. The studies presented herein, therefore, might be of crucial interest in understanding the etiology/pathology of MSUD in association with amyloid diseases and may pave for design of possible therapeutic remedies against the disease.

**Figure 1.**
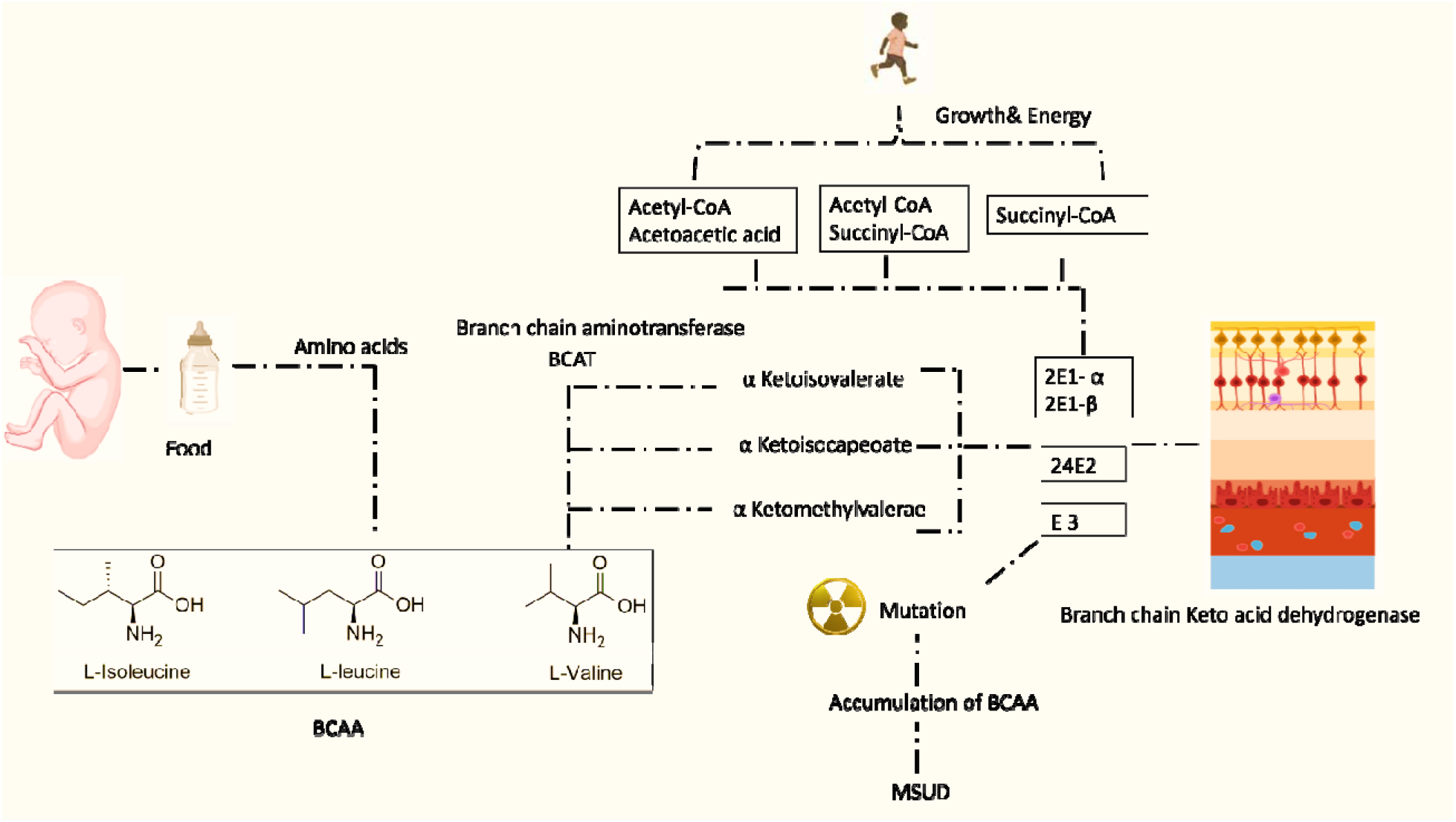
Schematic representation of accumulation of BCAA caused by gene mutation and consequence which leads to rare IEM such as MSUD.

## Result and Discussions

The chemical structure of the branched chain amino acids (BCAA) Isoleucine, Leucine and Valine reveal the amino acids might have amphipathic behavior due to hydrophobic aliphatic side chain and –NH_2_ and –COOH group which may interact with water (Figure 2a-c). Hence the chemical structure demonstrates a natural tendency of these amino acids to assemble in water. Hence, the self-assembling properties of BCAA were assessed by dissolving BCAA in water under neutral conditions.

**Figure 2.**
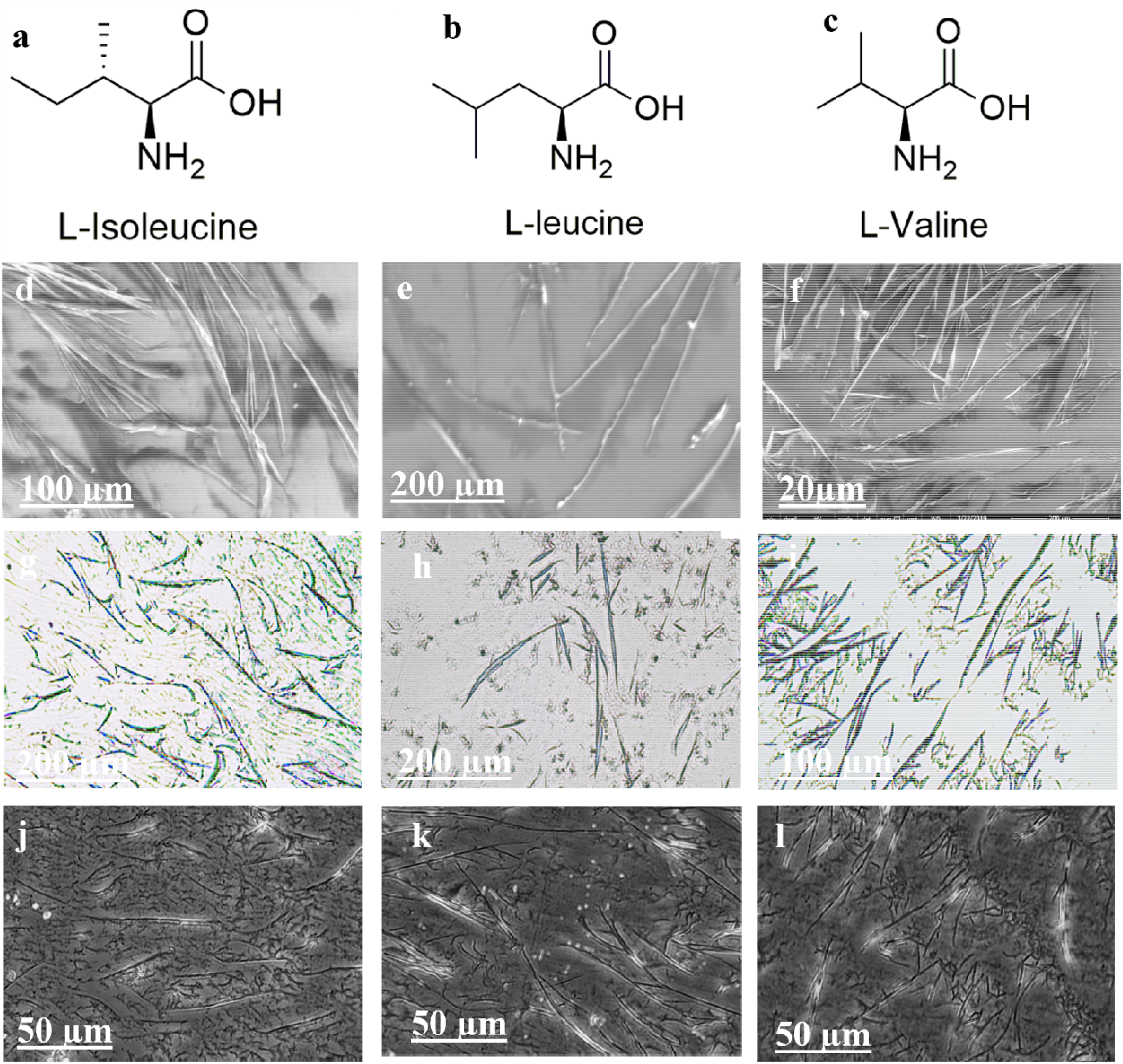
Chemical structures of BCAA (a) Ile; (b) Leu; (c) Val. Self-assembly of 1mM BCAA after 72 h in water, SEM images of (d) Ile; (e) Leu; (f) Val. Optical microscopy (OM) images in bright field mode (40X) of (g) Ile; (h)Leu; (i)Val. OM images in phase contrast mode (40X) of (j) Ile; (k) Leu; (l) Val.

Interestingly, 1mM solution of all the three branched chain amino acids assembled to febrile structure after 24-72 h of incubation. The fibers could be easily visualized by optical microscopy. Further, SEM was done to get insight into three dimensional febrile structures. Isoleucine, Leucine and Valine assembles to febrile structures with length ranging in several micrometers and diameter of fiber were varying in the range of 20-200 µm (Figure 2d-f). Another interesting observation was that in the SEM, BCAA assemblies tended to melt down as voltage or exposure time of the e-beam was increased. This observation further supports the conclusion that BCAA structures are indeed soft in nature and are formed exclusively via a self-assembly process. The melting might occur due to the release of trace water present in assemblies.^27^

To understand the role of intermolecular interactions in the self-assembly, we resorted to a series of comparative analyses of non-assembled and self-assembled Ile, Leu and Val using solid state NMR, FTIR, XRD and TGA.

### FTIR Interpretation of BCAA

To understand the role of hydrogen bonding and active involvement of functional groups in BCAA assembly process, FTIR spectra of Isoleucine, Leucine and Valine in self-assembled and non-assembled states were recorded. The aim was to characterize and investigate the difference in the behavior of amino acid molecules in assembled and non-assembled states through the variation in the vibration frequencies of constituent functional group. In this context, two main observations were deduced from the recorded FTIR spectra. Firstly, the observation that self-assembled state of amino acids exhibiting lower peak intensity compared to the non self-assembled state points towards the relative lowering of polarities around molecules primarily due to arrangement of molecules in a well packed manner after self-assembly.^28-31^ Secondly, the observed frequency shift is negligible in self-assembled single amino acid FTIR spectra in comparison to that of powdered amino acid sample. It clearly indicates that the covalent framework of the amino acid molecules during self-assembly of amino acid remains intact. Thus, self-assembly process is largely driven by only non-covalent forces such as van der Waals, π-π stacking, hydrogen bonding and other electrostatic interactions.^32^

Although, FTIR analysis did not yield any significant shifts or notable molecular change, data is indicative of new intermolecular interactions with BCAA carboxylic group. Quantitative analysis of peaks at 1500 and 3100 cm^-1^ show some (kubelka-monk method) showed some broadening by 5% an 8% respectively but it must be accompanied after NMR results. Non-assembled Ile showed two strong peaks at 1575.59, 1511.79 cm^-1^ peaks assigned to symmetric and anti-symmetric carboxylic group vibrations. A little change to 1575.42 cm^-1^ was detected after self-assembly (Figure 3a). Similarly, peaks at 2960.95 cm^-1^ assigned to CH stretching vibrations in non-assembled Ile are shifted slightly to 2961.75 cm^-1^. Hydroxyl stretching at 3100 cm^-1^ accompanied by 920.87 cm^-1^ out of plane bending vibration showed significant peak broadening while out of plane bending vibration intact at 920.8 cm^-1^, which can be attributed to the involvement of carboxylic acid group in temporary molecular interactions like hydrogen bonds resulting from self-assembly. Peak shifts in Leu were also quite comparable with Ile. Symmetric carboxylic group vibration in non-assembled Leu found at 15767.91 cm^-1^ was shifted to lower wavenumber 1577.6 cm^-1^ showing formation of hydrogen bond after self-assembly. CH stretching peaks in non-assembled Leu at 2618.8 cm^-1^ in are shifted to 2619.44 cm^-1^. Peak broadening effect is shown in hydroxyl peak at 3120cm^-1^. and 921.3 cm^-1^. These observations show carboxylic acid group in Leu is actively involving in self-assembly process (Figure 3b). Val also showed similar nature in FTIR spectra compared to Ile and Leu. Carbonyl stretching in non-assembled Val at 1563.31 cm^-1^ was shifted to 1565.12 cm^-1^. CH stretching at 2935.96 cm^-1^ is shifted to 2936.31 cm^-1^ after self-assembly. Peak broadening nature was also observed in hydroxyl stretching peak at 3100 cm^-1^ and 900.53 cm^-1^(Figure 3c). These observations show carboxylic acid group triggered the self-assembly process. In addition to that, a slight shift in NH deformation peak at 1504.72 cm^-1^ is shifted to 1505.73 cm^-1^ indicating the involvement of NH peak in self-assembly.

**Figure 3.**
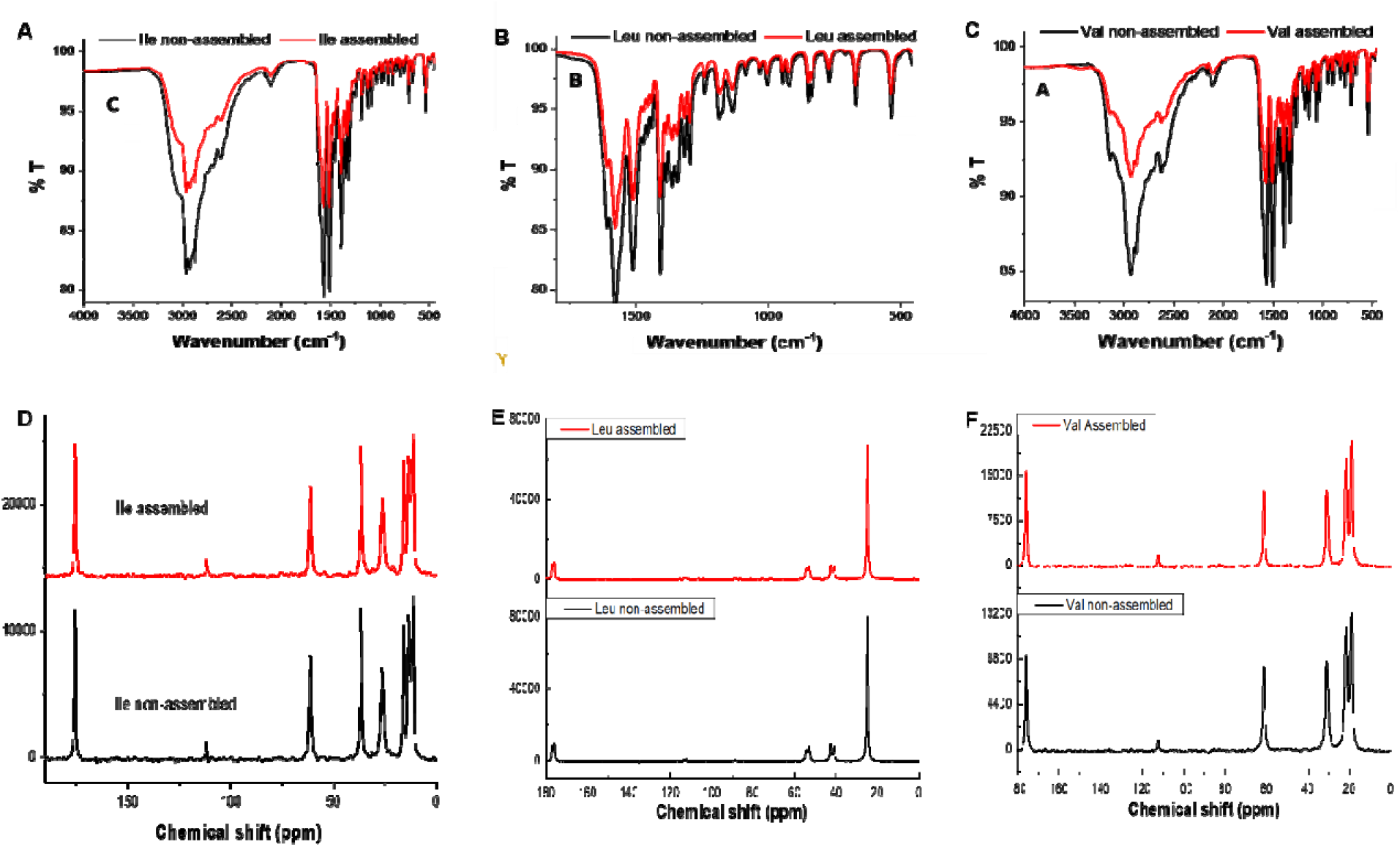
FTIR analysis of assembled and non-assembled BCAA. (A) Ile; (B) Leu; (C) Val. Solid-state NMR of assembled and non-assembled BCAA (D) Ile; (E) Leu; (F) Val.

Further, to verify the active involvement of carboxylic acid group, self-assembled solids of Ile, Leu and Val were isolated from 1 mM, 3 mM and 6 mM aqueous solution then analysed by ATR-FTIR in the same manner. In all cases noticeable peak broadening is observed at representative peaks of carbonyl and hydroxyl stretching after self-assembly (FigureS-1). Peak broadening effect was observed near carbonyl regions with increasing concentration from 1 mM to 9 mM. Interestingly intensity of the peaks around 1560 cm^-1^ and 900-930 cm^-1^ were decreased prominently showing effective intermolecular interactions. In case of Val, NH deformation peak at 1504.72 cm^-1^ also showed broadening effect upon increasing concentration to 9 mM indicative of hydrogen bonding nature. Overall transmission also gradually reduced with more concentration showing involvement of intermolecular interactions. Hence concentration dependent FTIR study indeed confirm carboxylic acid moiety of BCAA contributed more towards self-assembly and intramolecular interactions arising from polar ends like carboxylic acid groups in Ile, Leu and Val may play a significant role in self-assembly.

### Solid state NMR Interpretation of BCAA

Solid-state NMR spectra was used to investigate the molecular interactions and change in polymorphic nature after self-assembly. Non-assembled Ile shows multiplet peaks from 11-16 ppm for C6, C7 carbons. C8 peak appeared at 26.68 ppm with doublet nature and C5 singlet peak at 36.74 ppm. C4 carbon adjacent to polar amine group showed a sharp peak at 61.33 ppm and carbonyl carbon (C1) peak appeared at 175.56 ppm. After self-assembly, there is no significant peaks shift. However, side chain carbons exhibited lower chemical shifts at the same time carbonyl carbon (C1) and carbon adjacent to amine (C4) showed deshielding. These effects were indicative of intermolecular interactions involving carbonyl and amine groups and shows that side chain of Ile might be involved in the formation of an ordered structure. Significant peak broadening is observed particularly for carbonyl and C4 carbons and it confirm that molecular packing is changed due to self-assembly. C1 carbon peak position indicates there is no secondary structure formation but randomly ordered structures might be possible which is confirmed by microscopic images. It also indicates partial nature of COO^-^ ionisation (fully ionised C1 peak appears at 180 ppm and COOH at 167 ppm approximately) could be attributed to the formation of non-covalent bonds like hydrogen bonding at C1 carbon (Figure 3d).

Non-assembled Leu showed fully resolved peaks at 25.00 ppm for side chain C6, C7, C8 carbons and C5 carbon showed doublet at 41.02 ppm. C4 carbon peak adjacent to amine group appeared at 53.29 as doublet and carbonyl carbon (C1) appeared at 176.09 as a doublet. After self-assembly C5 proton exhibits slight shielding (up field) may be due to the rotation in C-C bond rotation caused by structural rearrangements. C4 proton showed slight de-shielding effect causing increased chemical shift due to new hydrogen bond formation with the amine group which reduces the effect of amine lone pair electrons on C4 carbon. C1 carbon in Leu exhibits noticeable downfield shift which confirms formation of a new hydrogen bond which lowers the electron density on C1 carbon as like Ile. Leu also shows peak broadening effect on C1, C4 and C6 carbons by 2-4 Hz indicating significant changes in polymorph after self-assembly induced by changes in the crystalline nature (Figure 3e).

Val showed separate singlet peaks for C6, C7 carbons at 21.54, 19.27 ppm due to repulsions occurring from carbonyl group. C5 and C4 carbons shows peaks at 31.12, 61.42 ppm respectively. Carbonyl carbon (C1) peak appears at 175.77 ppm after self-assembly. C5, C6, C7 carbons showed random shifts indicating significant changes in molecular arrangement possibly from C4-C5 single bond rotation. This bond rotation could be attributed to the non-covalent interactions of carbonyl oxygen with adjacent molecules induced by self-assembly. Since amide bond is planar and short side chain of Val may be forced the single bond rotation in order to create ordered structures after self-assembly (Figure 3f). Ordered self-assembled structures can be observed clearly by microscopic images. Significant peak broadening up to 4Hz (more than Ile and Leu) confirms major changes in morphology and molecular packing.

Solid-state NMR hence quietly reveals the involvement of carbonyl and amine groups in non-covalent interactions followed by lattice adjustment of side chain methyl groups responsible for self-assembly. It also reveals the absence of possible secondary structures. Placement of carbonyl peak reveals partial ionisation state indicative of new intermolecular interaction after self-assembly. Performing sophisticated CP-MAS NMR studies on each atom could be helpful to get detailed insights into molecular interactions after self-assembly which will be considered in our future work.

### Thermogravimetric analysis interpretation of BCAA

Thermogravimetric analysis (TGA) of Ile, Leu and Val was performed to gain insights of thermal stability and effect of molecular interactions after self-assembly (Figure 4). Generally branched chain amino acids show sharp melting ranges and sublimation effect on heating which is also observed in present study non-assembled Ile exhibited sharp and complete weight loss from 175 to 248 °C, indicating melting or sublimation effect. Self-assembled Ile exhibited weight loss of 96.8% from 161 to 235 °C. Self-assembly process resulted extra thermal stability owing to newly formed hydrogen bonds. Non-assembled Leu exhibited two weight loss segments first 4.97% below 150 °C and complete weight loss from 190 to 271 °C. Self-assembled Leu only one weight loss segment is shown from 174 to 258 °C. weight loss below 150 °C is due to loss of moisture attributed to high solubility of Leu in water. after self-assembly water loss peak is not significant. after self-assembly Leu observed slow degradation pattern and the unusual weight gain after 260 °C is also disappeared. This indicative of complete morphology changes after self-assembly induced by mew molecular interactions. Non-assembled Val showed complete weight loss from 195 to 274 °C while after self-assembly two weight loss regions are observed. first, weight loss of 2.1% below 158 °C due to moisture loss which could be trapped in pours structures formed after self-assembly. Second weight loss region observed from 174 to 256 °C with 99.7% weight loss. After self-assembly Val showed extra thermal stability and slow degradation rate which could be resulting from new molecular interactions. Similar observations can be seen clearly in DTG graphs which showing prominent shift in endothermic peaks in all the cases. DTG graph did not showed any unusual or new degradation path way after self-assembly concludes there is no molecular degradation or chemical occurred but morphological and physical property changes indeed took place. TGA analysis depicts that in all cases (Ile, Leu and Val) thermal degradation rate is decreased after self-assembly indicating presence of inter molecular interactions like hydrogen bonds and complete change in morphology/ polymorphic nature as observed by microscopy.

**Figure 4.**
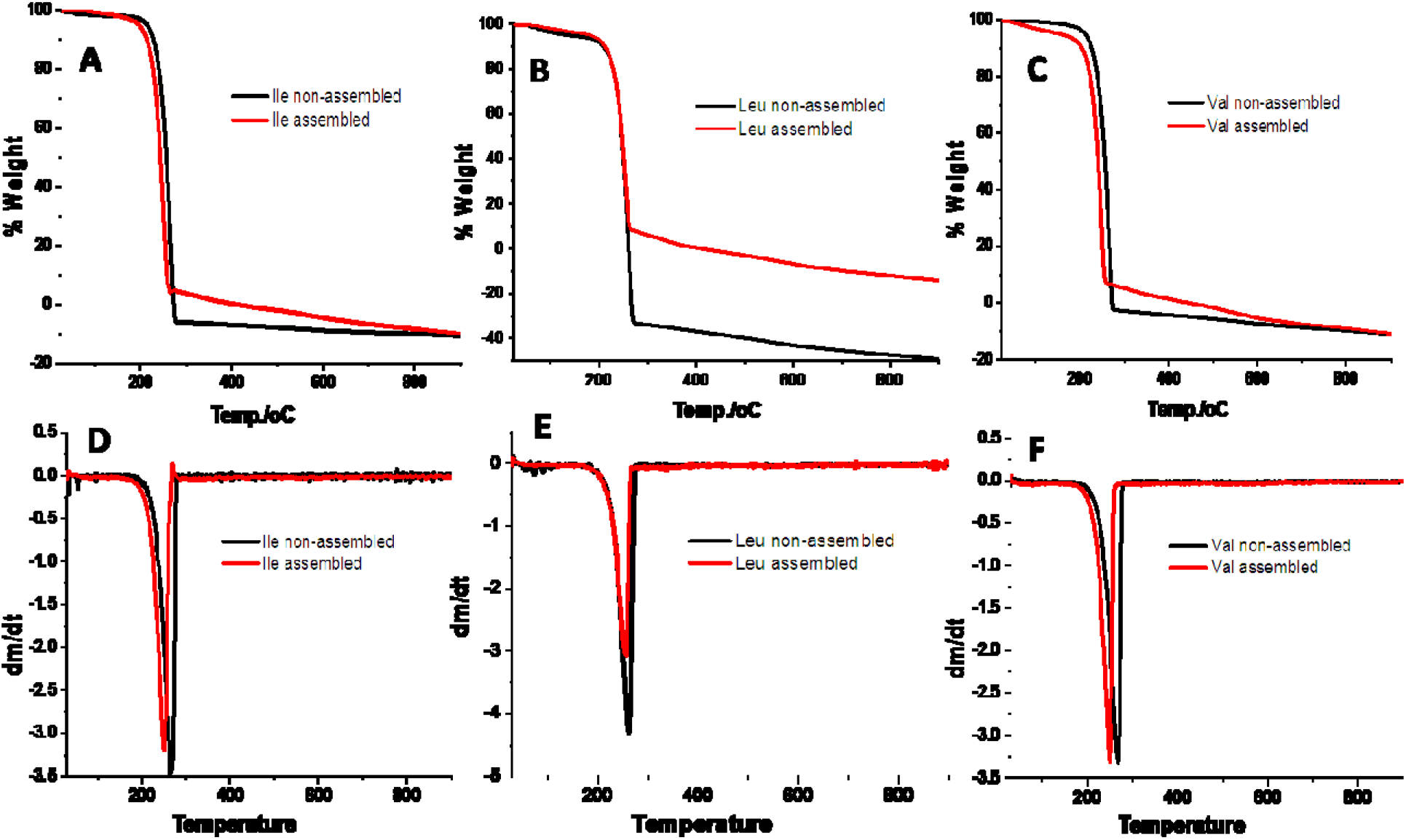
TGA and differential scanning calorimetry (DSC) analysis of assembled and non-assembled BCAA.

### XRD Interpretation of BCAA

Next, X-ray diffraction studies were done to investigate the crystalline nature and polymorphism in self-assembled amino acids. X-ray diffractograms showed some distinct characteristics like peak broadening, decrease in peak intensity and new peaks indicating increased polycrystalline or amorphous nature and change in molecular packing after the self-assembly process.

Ile non-assembled exhibited sharp peaks noticeably at 2θ 13.80 (88.47%), 20.17 (16.55%), 33.12 (77.01%), 39.70 (13.18%), 39.88 (11.42%), 46.54 (3.05%) and 74.54 (2.58%) indicating definite crystalline nature of the material. After self-assembly, increase in the relative intensity of diffraction peaks with slight shifts were observed. Peaks at 2θ 13.76 (96.63%) 20.16 (19.47%), 39.77 (14.86%), 74.54 (15.12%) revealed that peaks were broadened and intensity was decreasing suggesting change in molecular packing resulting from decrease in size and increase in density of crystalline aggregates (Figure 5a). Further, decrease in the peaks intensities at 2θ 33.12 (72.83%), 39.88 (0.12%) confirms the change in crystal packing nature. The peak broadening at 2θ 74.54 was significant and particularly indicates the presence of newly formed molecular interactions contributed by non-covalent forces after the self-assembly.

**Figure 5.**
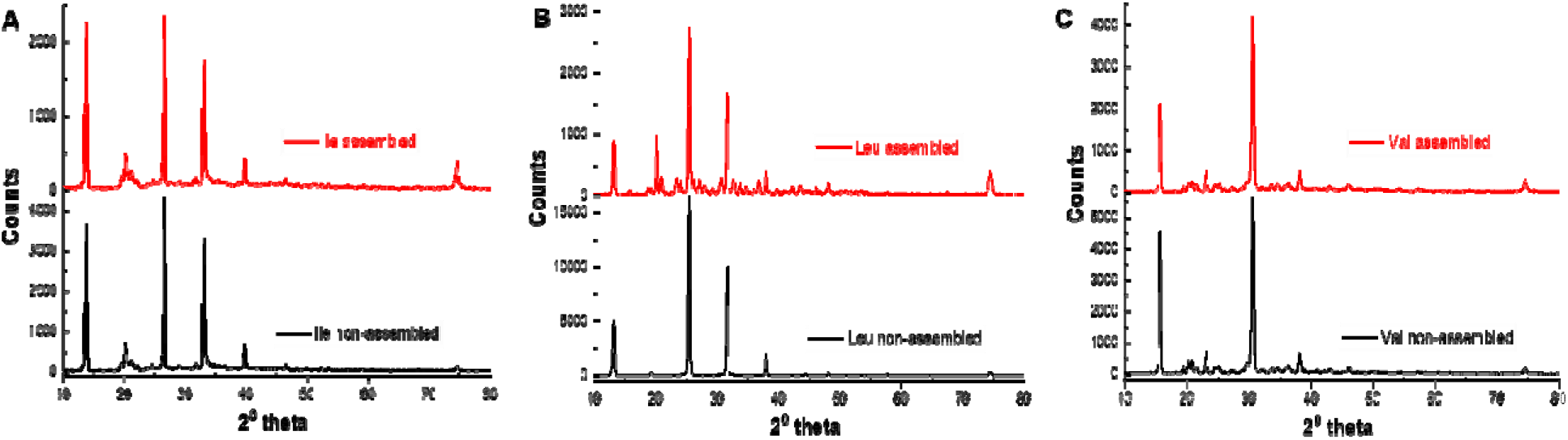
XRD analysis of BCAA before and after self-assembly. (A) Ile; (B) Leu; (C) Val.

Leu on the other hand, exhibited unusual diffraction patterns after self-assembly with drastic lowering of the peak intensity. This observation clearly changes in molecular packing and. formation of soft structures via self-assembly process. In case of Leu too the position of diffraction peaks remains unchanged but emergence of new diffraction peaks indicates formation of new morphologies with different lattice parameter Leu showed major diffraction peaks at 2θ 13.18 (30.40%), 25.43 (100%), 31.67 (61.16%), 37.98 (12.08%) and 74.52 (1.80%) indicating specific crystalline nature. After self-assembly Leu exhibited slight shifts with similar intensity whereas intensity of the peak at 2θ 74.48 increased to 13.06% with broadening indicative of strong π-π stacking (Figure 5b). Self-assembled Leu showed number of sharp peaks with relative intensity of 5-10% showing polycrystalline nature.

Similarly, Val also exhibited changes in diffraction pattern after self-assembly as like leu. Val non-assembled exhibited major diffraction peaks at 2θ 15.71 (80.65%), 19.42 (4,01%), 20.24 (6.27%), 23.11 (12.33%), 30.59 (100%), 38.21 (10.9%), 74.55 (3.10%) and many low intense sharp peaks indicating the crystalline nature of the material. After self-assembly, Val showed decrease in overall intensity of peaks and exhibited changes in the relative intensity of the diffraction peaks. Self-assembled Val showed relatively low intense peaks with slight shifts at 2θ 15.72 (50.08%), 19.45 (2.76%), 20.25 (4.49%), 23.11 (10.85%), 30.57 (100%), 38.21 (10.60%), 74.41 (6.08%) (Figure 5c). The XRD pattern in Val thus, indicates that self-assembled Val mostly retains a similar crystalline structure to that of non-assembled Val. However, emergence of new peaks in assembled Val indicates mixed polymorphic nature or formation of new crystalline forms after self-assembly.

Overall decrease in the peak intensity of Ile, Leu, Val in XRD studies confirmed that texture of the material was drastically changed after self-assembly.^32^ Peak broadening effect in most of the peaks indicative of smaller crystallite size with micro strain and reduced grain size. Particularly, decisive broadening effect was seen around 2θ 74° confirming the presence of newly formed ordered structures formed by non-covalent forces. The XRD studies clearly surmise that self-assembly changed lattice arrangements and induced polycrystalline to amorphous nature in BCAA. Peak shifts in all cases confirmed that self-assembly induces macro strain on the lattice which could be attributed to the formation of large aggregates. These random lattice arrangements can be confirmed by microscopic images. The non-crystalline nature of assemblies if further supported by phase contrast microscopic images.

In our previous studies, we have reported the crucial role of hydrogen bonding in single amino acid assembly of cysteine and methionine by co-incubation with urea.^22^ Since, urea breaks hydrogen bonds the assemblies would be disrupted if they are predominantly formed by hydrogen bonding. Interestingly when fibers of Ile, Leu and Val were co-incubated with urea, the assemblies were disrupted suggesting a crucial role of hydrogen bonding in BCAA aggregation (Figure 6).

**Figure 6.**
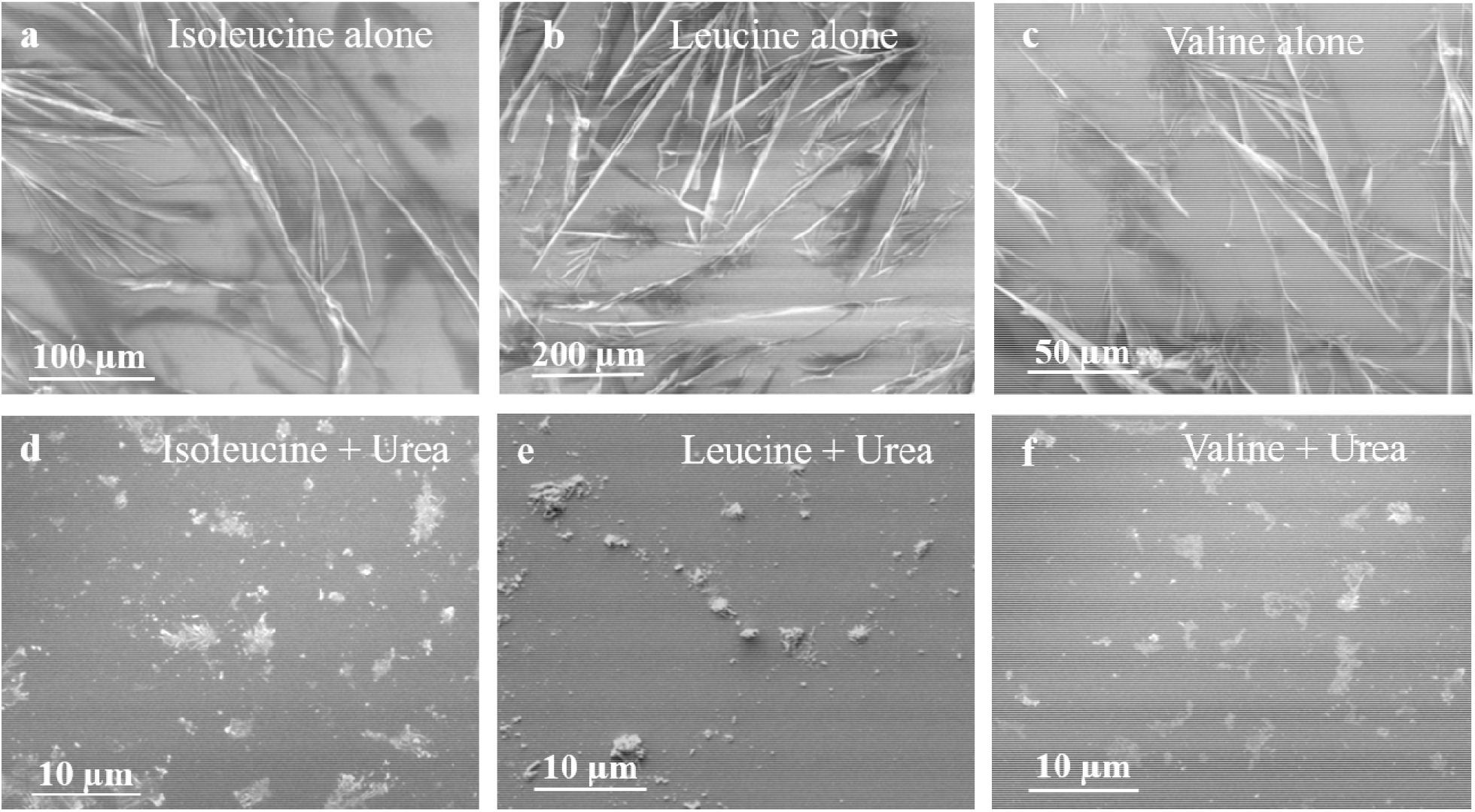
Representative SEM image of BCAA before and after co-incubation with Urea.

Additionally, to understand the role of water in formation of BCAA self-assemblies, Ile Leu and Val were incubated in an aprotic solvent mixture comprised of 3% DMSO: DCM and THF (Figure S2, S3).^27^ Since these solvents are aprotic they will not act as hydrogen bond donors, unlike water. Interestingly, under these conditions, no assembly was observed. However, as the trace water was added to THF, the BCAA self-assembly ensued and formation of fibrillar morphologies could be seen (Figure S3). Hence, these studies provided crucial evidence regarding role of water molecules in the structure formations of BCAA and its crucial significance in intermolecular hydrogen bonding.

Self-assembly is a spontaneous process of organization of molecules into well-ordered structures mainly driven by electrostatic interactions, hydrophobic attraction, hydrogen bonding and π-π stacking interactions.^34-43^ The results obtained via FTIR, solid state NMR,^38-41^ TGA and XRD of Ile, Leu and Val along with control experiments done using varying solvents and urea. suggest the crucial role of intermolecular hydrogen bonding in the self-assembly of both BCAA. Non-polar amino acids, like Ile, Leu and Val may assemble into fiber-like aggregates by hydrophobic interactions and possibly via backbone hydrogen bonding between the amine nitrogen and the carboxylic acid hydrogen present in an adjoining amino acid molecule.^34^ In addition, these molecules will have a tendency to form hydrogen bonds with polar solvents like water. The role of water molecules in formation of such self-assembled nanotubes/nanofibers has already been reported.^36,44^

With the fibers of Ile, Leu and Val being thoroughly characterized by solid state techniques like SEM, FTIR, TGA, NMR, we were next motivated to investigate the predominant solution structures of BCAA assemblies. Hence, ThT and CR binding assays were performed. ThT exhibits remarkable enhancement in its fluorescence on binding with amyloid.^45^ Fluorescence spectra of ThT with BCAA indeed revealed fluorescence enhancement and confirmed amyloid like structure formation by these single amino acids (Figure 7a). Interestingly the fibers also got stained with Thioflavin T dye indicating amyloid like characteristics of the assemblies (Figure 7). Since ThT binds to the hydrophobic pockets of β sheets present in amyloid fibers, it further confirmed BCAA like amyloid fibers do exist in solution. The amyloid nature of the assemblies was further confirmed using CR binding assay in solution. Although the manner of binding of CR with amyloidogenic peptides is still under debate, it is universally accepted that regardless of the manner of binding, CR displays an increase in absorbance intensity and a red-shift on binding with amyloid.^46,47^ Spectra obtained for BCAA with CR (Figure 7b) revealed a slight red shift and an increase in absorbance. As a control, we co-incubated other single amino acids namely, Phe, Tyr and Trp and also the dipeptide FF which has characteristic amyloid properties. All the reported single amino acids and FF exhibited an increase in absorbance and a similar red shift with CR. These results suggest that CR binds to structures formed by Cys and Met and hence the assemblies may have amyloid-like characteristics.

**Figure 7.**
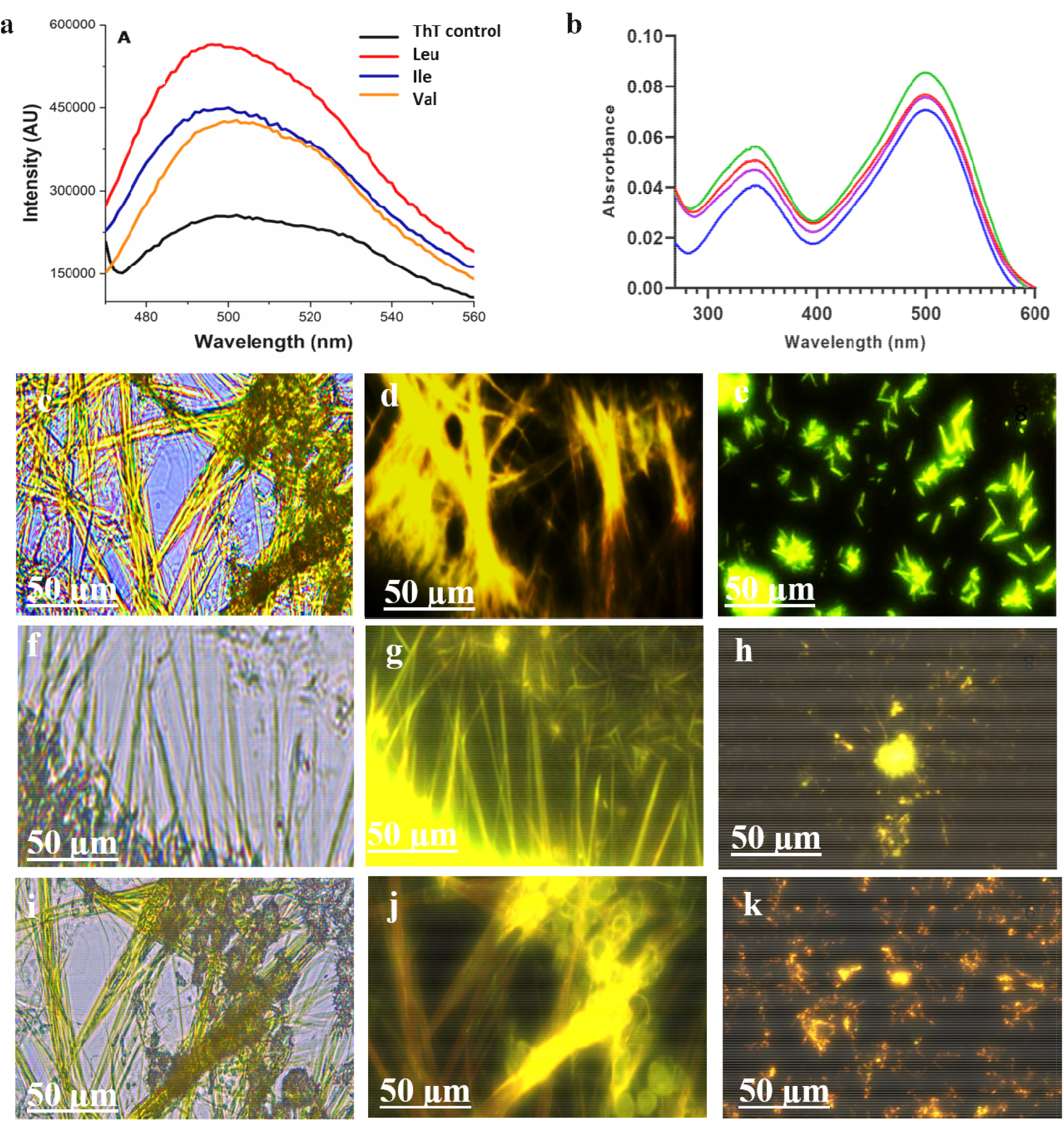
Amyloid-like properties of BCAA (a) ThT binding assay; (b) CR binding assay CR alone (blue), valine (violet), Isoleucine(red), Leucine (green). ThT stained microscopy images of 1 mM BCAAs (c, f, i) Ileu; (d, g, j) Leu; (e, h, k) Val.

### MTT assay of BCAA

Many amyloids are toxic as they tend to disrupt cell membrane, hence we were prompted to investigate the effect of these BCAA aggregates on cell viability. Therefore, to validate our hypotheses MTT assay of RPE-1 cells co-incubated with fresh and aged samples of BCAA assemblies were performed in buffered conditions. MTT assay is generally a quick colorimetric test that measures the number of live cells in living cells by measuring the cleavage of the tetrazolium ring of MTT (3-(4,5-dimethylthazolk-2-yl)-2,5-diphenyl tetrazolium bromide) by dehydrogenases in active mitochondria. The MTT assay suggests aged samples of Ile, Leu and Val caused drastic decrease in cellular viability in RPE-1 cell lines in PBS (100mM, pH 7.4) (Figure 8). The study confirms cytotoxic nature of aged assemblies. The MTT assay of Ile, Leu and Val assemblies after 3 days of ageing reveals dose dependent toxicity. The MTT assay thus confirmed the cytotoxic nature of these assemblies.

**Figure 8.**
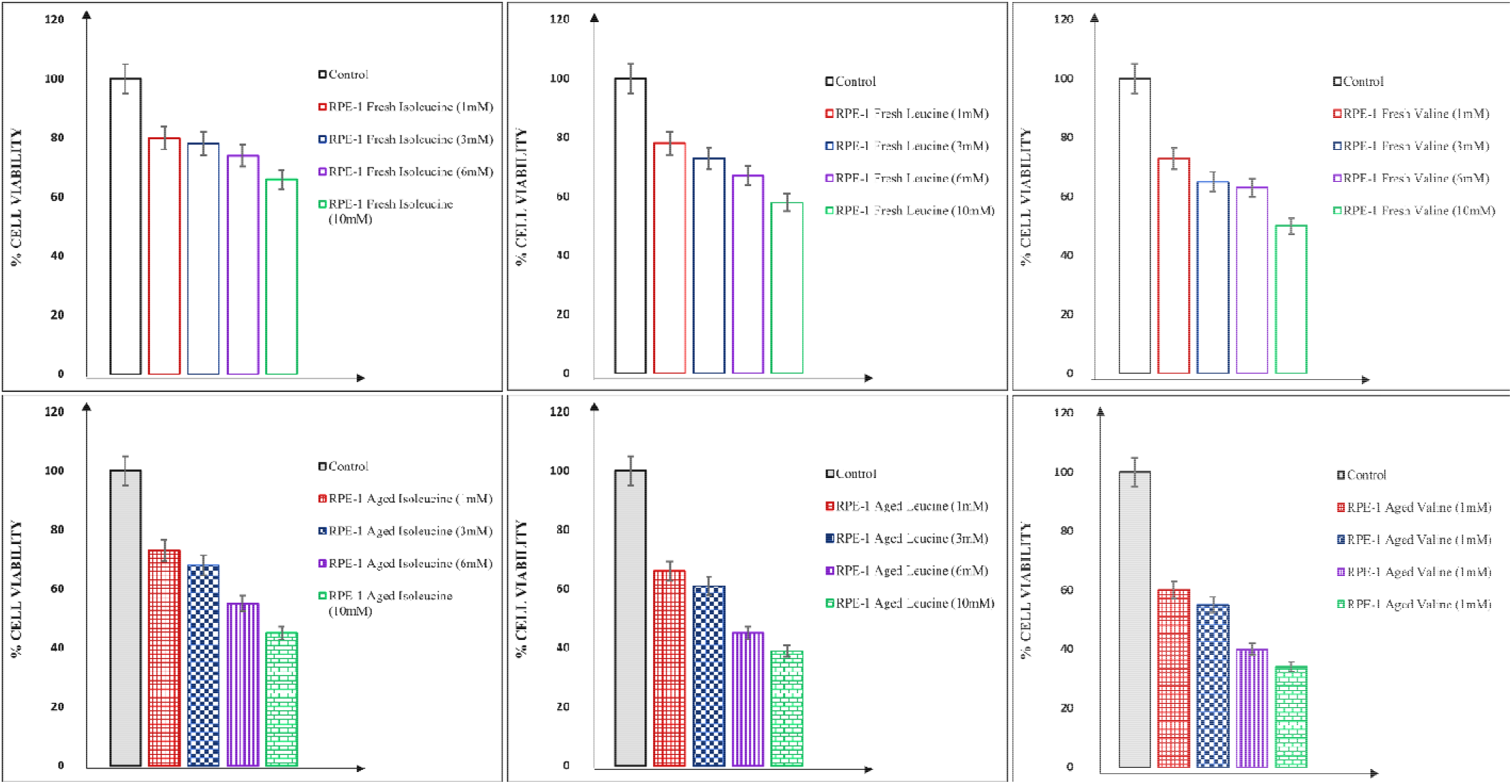
MTT assays after co-incubation with varying concentrations of Ile, Leu and Val in fresh and aged conditions in RPE-1 cell lines.

## Conclusions

In summary, we have investigated the self-assembly properties of BCAA through various niophysical assays and microscopy studies. The intermolecular forces which causes amyloid-like structure formation were probed through solid state NMR, FTIR, XRD and TGA. Since these structures bind amyloid specific dyes like ThT and CR and reveal cytotoxic nature as the cell viability was reduced in RPE-1 cell lines. Thus, Amyloid binding assays and cell viability analyses revealed that aggregates may have an amyloid nature and are cytotoxic. These results are thus highly significant and indicate physicochemical properties of MSUD might be related to toxic aggregation of BCAA and may have immense implication in understanding the pathogenesis of the disease.

### Experimental section

The metabolites used in this experiment were of analytical grade having purity <99% obtained from SRL and Avra chemicals and were used without further purification. The stock solution (100 mM) of the metabolites was prepared in DMSO and further diluted to their desired concentration using deionized water. These solutions were incubated for varied amount of time at 37.4C. Each sample was spread out over a glass slide in 20 µL portions for OM imaging, and then it dried. The images were taken using Leica DM2500 microscope under 40X and 63X magnifications.

### Optical Microscopy (OM)

The self-assembling characteristics of the samples were examined using OM by dropcasting a 20 µL solution at varying concentrations and ageing time, which was then dried at 37.4 □C on a clean glass slide. Leica DM2500 microscope with 40X and 63X magnifications were used to capture all the images.

### Scanning Electron Microscopy (SEM)

The SEM images were taken using a Nova Nano FEG-SEM 450 microscope, which was operated at an accelerating voltage of 5 to 15 kV. SEM samples were prepared using Silicon wafers. The samples were dispersed and dried at 37.4 □C.

### Coincubation assay with urea

For coincubation experiment, the BCAA were coincubated with urea in equal ratio (1:1). The final concentration of BCAA and urea was made to be similar. Both were mixed and incubated for three to four hours.

### ThT assay

A stock solution of 5 mM ThT was made in Milli-Q water for the ThT binding experiment. The equal concentration of BCAA and ThT dye was taken and coincubated for 30 minutes at 37.4 □C. The fluorescence spectra was recorded using Cary Eclipse Fluorescence Spectrophotometer. With a wavelength of excitation of 460 nm and an emission slit of 5 nm, all the data were captured.

### Congo red (CR) binding assay

For CR assay, the samples were prepared in deionized water and incubated for 12 h at 37.4 □C. The concentration of BCAA was 100 µL solution and was added 2 µL of CR (stock of CR prepared was 500 µM) and the final concentration of CR in solution was 10 µM. The UV-vis spectra of the saples was recorded using UV-1900 SHIMADZU spectrophotometer after 10 min of coincubation with CR.

### Fourier Transform Infrared (FTIR) Spectroscopy

FTIR analysis of the BCAAs were done in DMSO solvent using Shimadzu IR Affinity-1S spectrophotometer instrument. The spectra were recorded between 4000 cm^-1^ to 400 cm^-1^ in wavenumber. At different concentrations, the FTIR spectra of self-assembled BCAAs (lyophilized powder) were recorded and compared to those of the commercial available BCAAs. The self-assembled lyophilized powder of Ise, Leu and Val in deionized water were incubated for 12 h at 37.4 □C. The solution was dried using lyophilizer (Martin Christ) in order to prevent the assemblies from disrupting.

### Solid state ^1^H NMR Spectroscopy

Solid state ^1^H NMR of self-assembled and non-assembled samples of BCAAs were recorded on BRUKER 400 MHz instrument at 37.4C. The solid sample about 100 mg was placed in the sample holder and the spectra was recorded at 5 kHz energy.

### Thermogravimetric Analysis (TGA)

TGA studies of BCAAs were performed using a PerkinElmer TGA instrument (TGA-4000) under nitrogen atmosphere. Both the samples i.e. self-assembled and commercial BCAAs were placed in a ceramic crucible and heated from 300 to 900 °C at a fixed heating rate of 10 °C/min. The self-assembled lyophilized powder of BCAAs were also made similar to FTIR samples using 5 mM solutions in deionized water. The TGA spectra of this lyophilized powder and the powder of the BCAA that was bought commercially were then compared.

**X-ray Diffraction (XRD) Analysis:** Pan-analytical X-pert Pro instrument was used to perform XRD experiment. The samples were scanned within the range of 2θ 10−80° after being equally dispersed over the substrate holder. The samples for XRD were prepared by lyophilizing a 3 mM solution of BCAAs 12 h before lyophilization.

### *In vitro* cytotoxicity analysis

The cytotoxicity analysis on RPE-1 cell lines was performed via MTT assay by following previously reported methodologies. The cells were purchased from National Centre for Cell Science (NCCS), Pune, India. RPE-1 cells (2×105 cells/mL) were cultured in 96-well micro plates (100 mL/well) and incubated overnight at 37°C. 100 mL of amino acids was dissolved in Dulbecco’s Modified Eagle Medium (DMEM) at various concentrations was then thrown over cells. The samples were aged optimally for 3 days in DMEM media before incubating them with cells.

## Acknowledgements

NG greatly acknowledge support from SERB research grant SERB SPG/2021/000521 for funding and fellowships and Indrashil University for infrastructure support. DB thanks SERB-CRG, MoES-STARS, Gujcost-DST, GSBTM and IITGN for research grants. MKP would like to thank the Department of Science and Technology, Govt. of India research grant (DST/TM/WTI/WIC/2k17/100). The work was supported by the SERB SPG/2021/000521 received by Dr. Nidhi Gour.

## Supporting Information Available

The supporting information contains additional data regarding experiments and supplementary Figures which are referred as Figure S-1 - S-3 in the main text.

