## Supporting information for "Assembly of branched chain amino acid (BCAA) to toxic fibrils may be related to pathogenesis of Maple syrup urine disease (MSUD)"

| **Sr.No.** | **Topic** | **Page No.** |
| --- | --- | --- |
| 1. | Experimental procedures  Optical microscopic studies | 2 |
| 2. | Fourier Transform Infrared (FTIR) Spectroscopy | 2 |
| 3. | Solvent-dependent studies | 2 |
| 4. | Optical microscopy, SEM and FTIR studies | 3-4 |

**Experimental procedures:**

1. **Optical microscopic studies:**

The BCAAs used in this study were of analytical grade and purchased from SRL and Avra chemicals and used without further purification. The sample preparations were done in deionized water. The stock solution (100 mM) of the metabolites was prepared in DMSO and further diluted to their desired concentration using deionized water. These solutions were incubated for varied amount of time at 37.4 C. Each sample was spread out over a glass slide in 20 µL portions for OM imaging, and then it dried. The images were taken using Leica DM2500 microscope under 40X and 63X magnifications.

2. **Fourier Transform Infrared (FTIR) Spectroscopy:** FTIR analysis of the BCAAs were done in DMSO solvent using Shimadzu IR Affinity-1S spectrophotometer instrument. The spectra were recorded between 4000 cm^-1^ to 400 cm^-1^ in wavenumber. At different concentrations, the FTIR spectra of self-assembled BCAAs (lyophilized powder) were recorded and compared to those of the commercially available BCAAs. The self-assembled lyophilized powder of Ise, Leu and Val in deionized water were incubated for 12 h at 37.4 C. The solution was dried using lyophilizer (Martin Christ) in order to prevent the assemblies from disrupting.
3. **Solvent-dependent studies:** The solvent-dependent study was performed to understand the role of water in formation of BCAA self-assemblies. Herein, different solvents such as tetrahydrofuran (THF), dimethyl sulfoxide (DMSO) and dichloromethane (DCM) was used at optimal conditions. The samples of BCAA were incubated in DMSO:DCM and water:THF respectively, to conduct the self-assembly study of these compounds. The samples were incubated at 37.4 °C and 20 μL of each sample was spread on a clean glass slide for OM and left to dry. Once the samples were dried, images were captured using a Leica DM2500 microscope at 40X and 63X magnification under bright field.
4. **Optical, SEM and FTIR studies:**

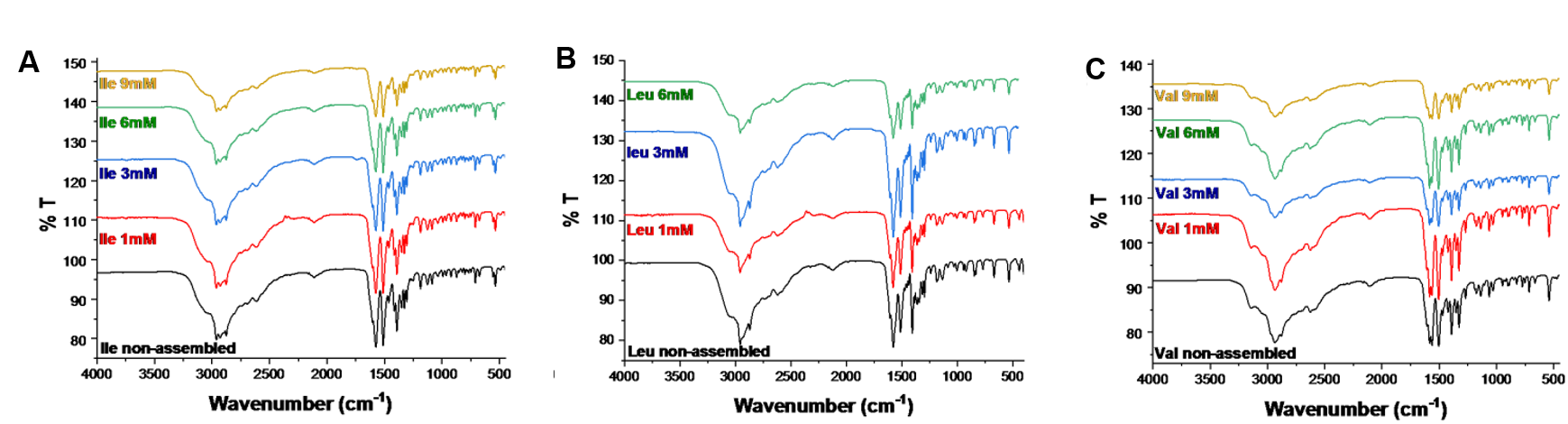

Figure 1: Concentration dependent FTIR analysis of (A) Ile; (B) Leu and (C) Val.

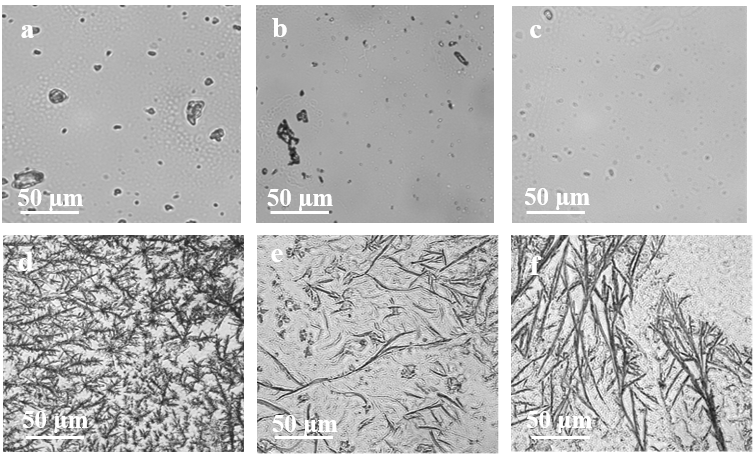

Figure 2: Optical microscopy image of BCAA in 3% DMSO:DCM (a) Ile; (b) Leu; (c) Val, show no fibrillar morphologies. SEM image of BCAA in 3% Water: THF shows regeneration of fibrillar morphologies; (d) Ile; (e) Leu; (f) Val, showing regeneration of fibrillar structures.

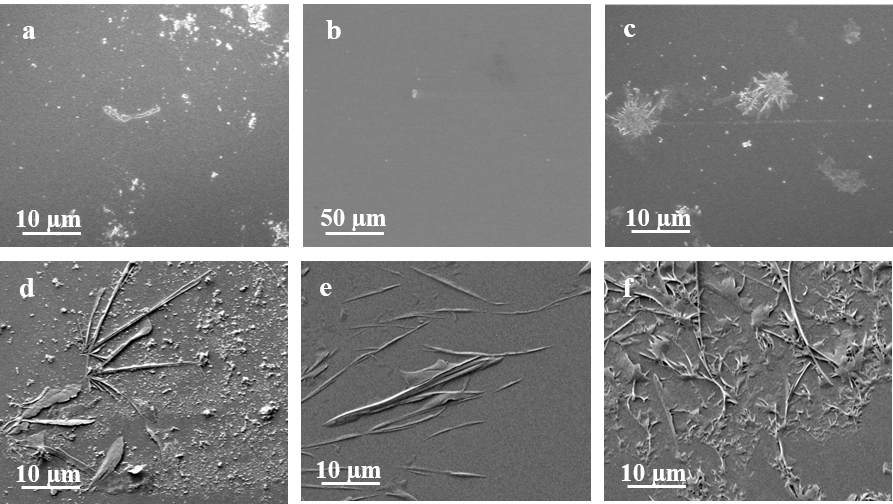

Figure 3: SEM image of BCAA in 1% Water: THF (a) Ile; (b) Leu; (c) Val. SEM image of BCAA in 3% Water: THF showing regeneration of assemblies (d) Ile; (e) Leu; (f) Val.
